# The WT1-like transcription factor Klumpfuss maintains lineage commitment in the intestine

**DOI:** 10.1101/590885

**Authors:** Jerome Korzelius, Tal Ronnen-Oron, Maik Baldauf, Elke Meier, Pedro Sousa-Victor, Heinrich Jasper

## Abstract

Stem cell (SC) lineages in barrier epithelia exhibit a high degree of plasticity. Mechanisms that govern the precise specification of SC daughter cells during regenerative episodes are therefore critical to maintain homeostasis. One such common mechanism is the transient activation of the Notch (N) signaling pathway. N controls the choice between absorptive and entero-endocrine cell fates in both the mammalian small intestine and the *Drosophila* midgut, yet how precisely N signaling promotes lineage restriction in progenitor cells remains unclear. Here, we describe a role for the WT1-like transcription factor Klumpfuss (Klu) in restricting the fate of *Drosophila* enteroblasts (EBs) downstream of N activation. Klu is transiently induced in Notch-positive EBs and its transient activity restricts cell fate towards the enterocyte (EC) lineage. Transcriptomics and DamID profiling show that Klu suppresses enteroendocrine (EE) cell fates by repressing E(Spl)m8-HLH and Phyllopod, both negative regulators of the proneural gene Scute, which is essential for EE differentiation. At the same time, Klu suppresses cell cycle genes, committing EBs to differentiation. Klu-mediated repression of its own transcription further sets up a negative feedback loop that ensures temporal restriction of Klu-mediated gene regulation, and is essential for subsequent differentiation of ECs. Our findings define a transient cell state in which EC lineage restriction is cemented, and establish a hierarchy of transcriptional programs critical in executing a differentiation program downstream of initial induction events governed by N signaling.

## Introduction

In many tissues, somatic SCs respond to tissue injury by increasing their proliferative potential and producing new differentiating cell progeny. To maintain homeostasis during such periods of regeneration, cell specification and differentiation need to be precisely coordinated within a dynamic environment. Studies in the mammalian intestine have demonstrated a surprising plasticity in such specification events, showing that even differentiated cells can revert into a stem cell state in conditions in which tissue homeostasis is perturbed ^1,2^. These findings highlight the critical role of gene regulatory networks in establishing and maintaining differentiated and committed cell states in homeostatic conditions.

The *Drosophila* midgut is an excellent model to study lineage differentiation of adult stem cells both in homeostasis as well as during regeneration and aging. The *Drosophila* midgut is maintained by intestinal stem cell (ISCs), which can generate differentiated enterocytes (EC) or entero-endocrine (EE) cells ^3,4^. Upon injury or infection, ISC proliferation is dramatically increased in response to mitogenic signals from damaged enterocytes ^5–7^. Mis-regulation of cell specification and differentiation in this lineage can lead to significant dysfunction, as evidenced in aging intestines, where disruption of normal N signaling due to elevated Jun-N-terminal Kinase (JNK) signaling leads to an accumulation of mis-differentiated cells that contribute to epithelial dysplasia and barrier dysfunction ^8,9^.

Notch signaling plays a central role in both ISC proliferation and lineage differentiation. ISCs produce the Notch-ligand Delta and activate Notch in the EB daughter cell. Levels of Delta vary markedly between ISCs in the homeostatic intestine. These differences have been proposed to underlie the decision between EC and EE differentiation in a specific ISC lineage ^10^. It has been proposed that higher N activity is associated with differentiation into the EC fate, while lower N activity promotes EE differentiation ^10,11^. Loss of N in ISC lineages leads to the formation of tumors that consist of highly Delta-expressing ISCs and of Prospero (Pros)-expressing EEs ^10,12,13^. These tumors are likely a consequence of impaired EB differentiation, resulting in an increased frequency of symmetric divisions, as well as of differentiation of a subset of EBs into EE, suggesting that EE differentiation is a default fate in the lineage when N signaling activity is reduced.

Interestingly, recent work has shown that lineage specification in ISC daughter cells is likely more complex than a simple model in which EB fates are determined stochastically or by ‘lateral-inhibition’ – like N-mediated mechanisms. It has been shown that pre-determined ISCs exist that express the EE marker Prospero and generate daughter cells that differentiate into EEs ^14,15^.

The exact cell state in which the decision between EE and EC fates is cemented, however, remains unclear. A transient specification step has been identified in EE differentiation, in which cells transiently express Scute, a transcription factor that negatively regulates N responsive genes such as E(Spl)m8, as well as its own expression ^16^. Furthermore, EBs have been shown to remain in a transient state for a prolonged period of time before differentiating into an EC fate ^17^. Mechanisms that regulate and maintain this transient state remain unclear.

Here we describe a role for the Wilms’ Tumour 1 (WT1) – like transcription factor Klumpfuss (Klu) in lineage commitment during EC differentiation in the adult fly intestine. In transcriptome studies of FACS sorted ISCs and EBs, we found Klu to be expressed in an Esg-dependent manner specifically in EBs. Klu is critical for leg and bristle development, as well as for lineage differentiation of Type II neuroblast stem cells ^18,19^. Klu is related to the mammalian tumor suppressor gene Wilms’ Tumour 1 (WT1), and its overexpression in neuroblast stem cells leads to tumorous overgrowths that can grow autonomously when transplanted in the abdomen of flies ^20,21^. In the intestine, we find that loss of Klu leads to aberrant entero-endocrine differentiation of EB cells, whereas ectopic activation of Klu results in a failure to differentiate. Transcriptomics and DNA-binding studies reveal that Klu controls EE differentiation by repressing genes involved in Notch signaling, as well as by indirectly controlling the levels of the Achaete-Scute complex members *asense* and *scute*. Klu further represses its own expression, defining a transient state of EBs in which specification into ECs is cemented by precise temporal regulation of N signaling. We propose that the transient expression of Klu ‘locks in’ the EC fate in EBs, preventing ectopic proneural gene activation and thus ensuring lineage commitment into the EC fate.

## Results

### The transcription factor Klu is expressed in the enteroblast precursor cells

We identified Klumpfuss (Klu) transcripts to be specifically enriched in transcriptomes isolated from sorted progenitor cells ^22^ and to be significantly downregulated upon loss of the stem and progenitor specific transcription factor Escargot (Esg) ^23^. To confirm and better characterize *klu* expression in the *Drosophila* posterior midgut, we used a *klu-Gal4, UAS-GFP* reporter line that reflects Klu expression in the wing and eye discs of wandering 3rd instar larvae^18,24^. In the midgut, GFP expression was seen in the larger cells of the stem-progenitor nests (ISC+EB). These cells resemble EBs based on both nuclear and cellular size (Figure 1A arrowheads). To confirm their identity, we combined the Klu reporter line with the Notch activity reporter *Su(H)GBE-lacZ*, which is exclusively activated in EBs, but not in ISCs ^10^. In addition, we used *Delta-lacZ* (*Dl-lacZ*) as a marker for ISCs. The expression of *klu-Gal4, UAS-GFP* overlapped almost exclusively with the EB-specific *Su(H)GBE-lacZ* reporter. In contrast, ISC-specific Delta-lacZ staining was mostly found in small, diploid cells neighboring the GFP-positive cells (Figure 1 B, C, quantification in D, E).

**Figure 1.**
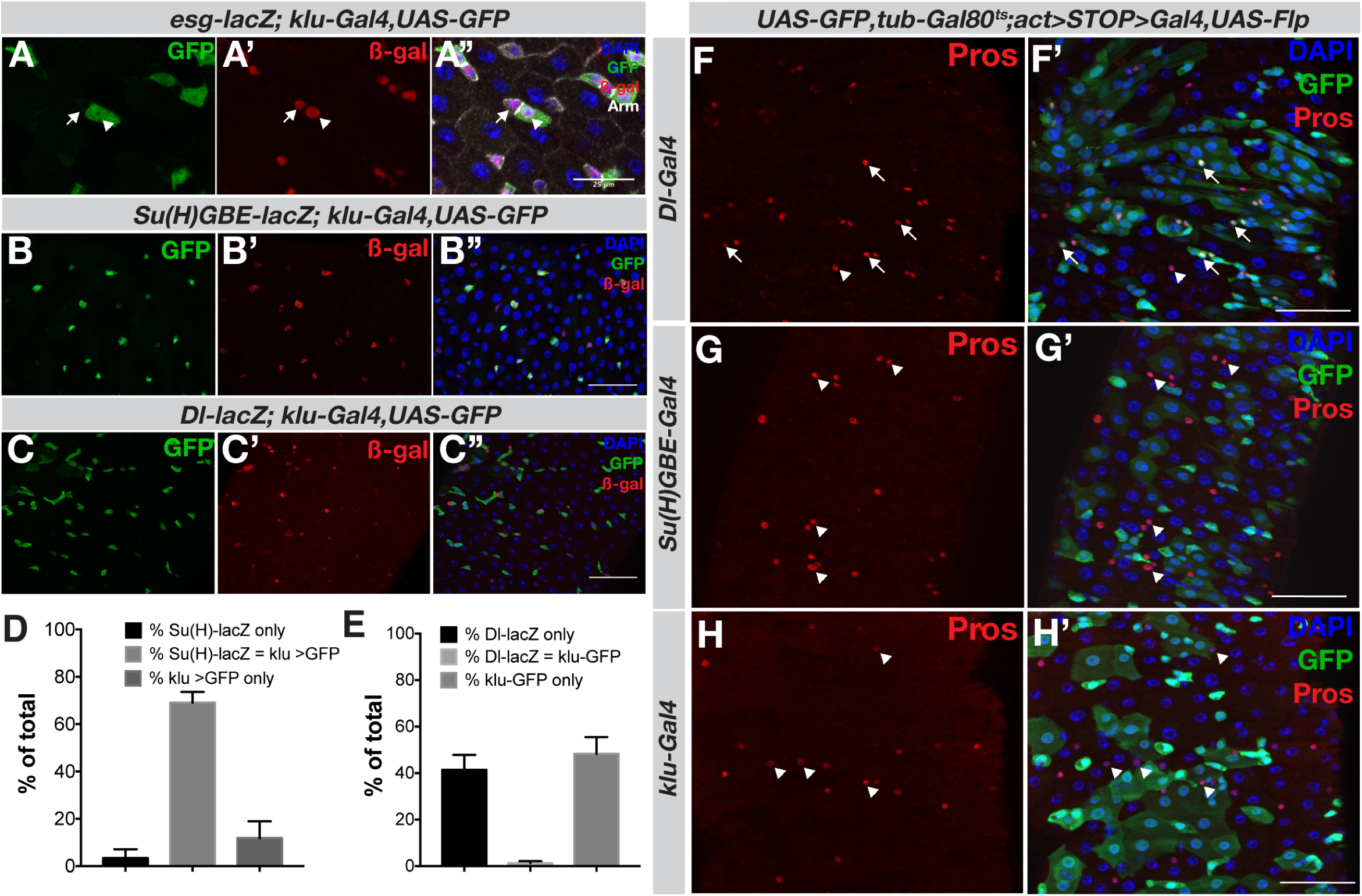
The transcription factor Klu is specifically expressed in enteroblast cells. **A-A’** The *klu-Gal4, UAS-GFP* reporter line shows expression in the midgut epithelium. ISCs (arrows) and EBs (arrowheads) are visualized by *esg-lacZ*. Cells are outlined with Armadillo/β-catenin. **B-C**. The *klu-Gal4, UAS-GFP* was combined with *Su(H)-GBE-lacZ* (enteroblast (EB) marker) or *Dl-lacZ* (intestinal stem cell (ISC) marker). Expression of Klu largely overlaps with the EB marker *Su(H)-GBE-lacZ* (B), and Klu-positive cells are found adjacent to the Delta-positive ISC (C). **D-E**. Quantification of marker gene overlap of the genotypes displayed in (**B-C**) *n* = 7 guts (*Dl-lacZ, n* = 1370 cells counted) and *n* = 10 guts (*Su(H)-GBE-lacZ, n* = 572 cells counted). **F-H**. Lineage-tracing of cells in the intestine using different cell-specific drivers. **F-F’**. The *Dl-Gal4*-positive ISCs give rise to both differentiated cell types of the intestinal lineage (enterocytes (EC) and entero-endocrine (EE) cells). EEs are marked by antibody staining for the transcription factor Prospero (Pros, in red). Arrows indicate GFP-Pros double-positive EEs in the clonal area, whereas arrowheads indicate EEs outside the clonal area. **G-G’**. Su(H)-GBE-positive EB cells exclusively give rise to ECs, but not to EEs (arrowheads). **H-H’**. Similar to Su(H)-GBE-positive EBs, Klu-positive cells also give rise exclusively to ECs, but not EEs.

Lineage tracing experiments have previously shown that Notch-positive EB precursor cells exclusively give rise to enterocytes, whereas Delta-positive ISCs can give rise to clones with both ECs and EEs ^14,15^. To trace the fate of Klu-expressing cells, we crossed the klu-Gal4 enhancer-trap line to a Actin promoter-driven FlipOut lineage tracing cassette (*UAS-GFP, tub-Gal80ts;UAS-Flp, Act>STOP>Gal4*). Upon temperature shift, the UAS-Flp-inducible *Act>STOP>Gal4* will be activated, marking all cells that express Klu as well as all differentiated progeny from these cells. As expected, Dl-Gal4 expressing ISCs give rise to both ECs (visible as large GFP-positive cells with a large, polyploid nucleus) as well as EEs, marked by expression of the transcription factor Prospero (Pros) (Figure 1F, arrows). In contrast, Notch-positive EBs (Su(H)GBE-Gal4) only gave rise to ECs, but not EEs (Figure 1G, arrowheads). Similar to Notch-positive EBs, *klu-Gal4*-traced cells gave rise exclusively to ECs, but not EEs (Figure 1H). We conclude that Klu is specifically expressed in the EC-generating EBs in the *Drosophila* midgut.

### Klu loss of function leads to excess EE differentiation

To determine the role of Klu in the specification and/or differentiation of cells in the ISC lineage, we first inhibited Klu function using RNAi. We used the TARGET-system to express UAS-driven RNAi constructs in specific lineages in a temperature-dependent manner ^25^. We used the *esg-Gal4^ts^* driver to express Klu RNAi in ISCs and EBs and the *Su(H)GBE-Gal4^ts^* driver that drives expression in EBs only. In both conditions, knockdown of Klu lead to an increase of EEs in the posterior midgut (Figure 2A-D, quantification in E). To confirm the EE differentiation phenotype, we used Mosaic analysis with a repressible cell marker (MARCM) ^26^ to generate marked lineages homozygous for a null allele of Klu, *klu^R51^* ^18^ and traced the fate of *klu^R51^* mutant cells. Quantification showed that loss of Klu leads to more EE cells/clone (Figure 2F-I). Interestingly, the GFP-negative tissue also contained more EEs in *klu^R51^* MARCM animals than in the control animals (*FRT2A*, Figure 2F, compare with 2G). This is likely due to the fact that in this genotype, the GFP-negative tissue is heterozygous for the *klu^R51^* null allele. In addition, we generated stem cell clones expressing *klu^RNAi^* using the *escargot* promoter-driven FlipOut system (*esg-F/O*). Enterocyte differentiation still occurred in *esg-F/O > klu^RNAi^* clones based on nuclear size of cells within the clones and staining for the enterocyte marker Pdm1 ^23,27^ (Figure S1).

**Figure 2.**
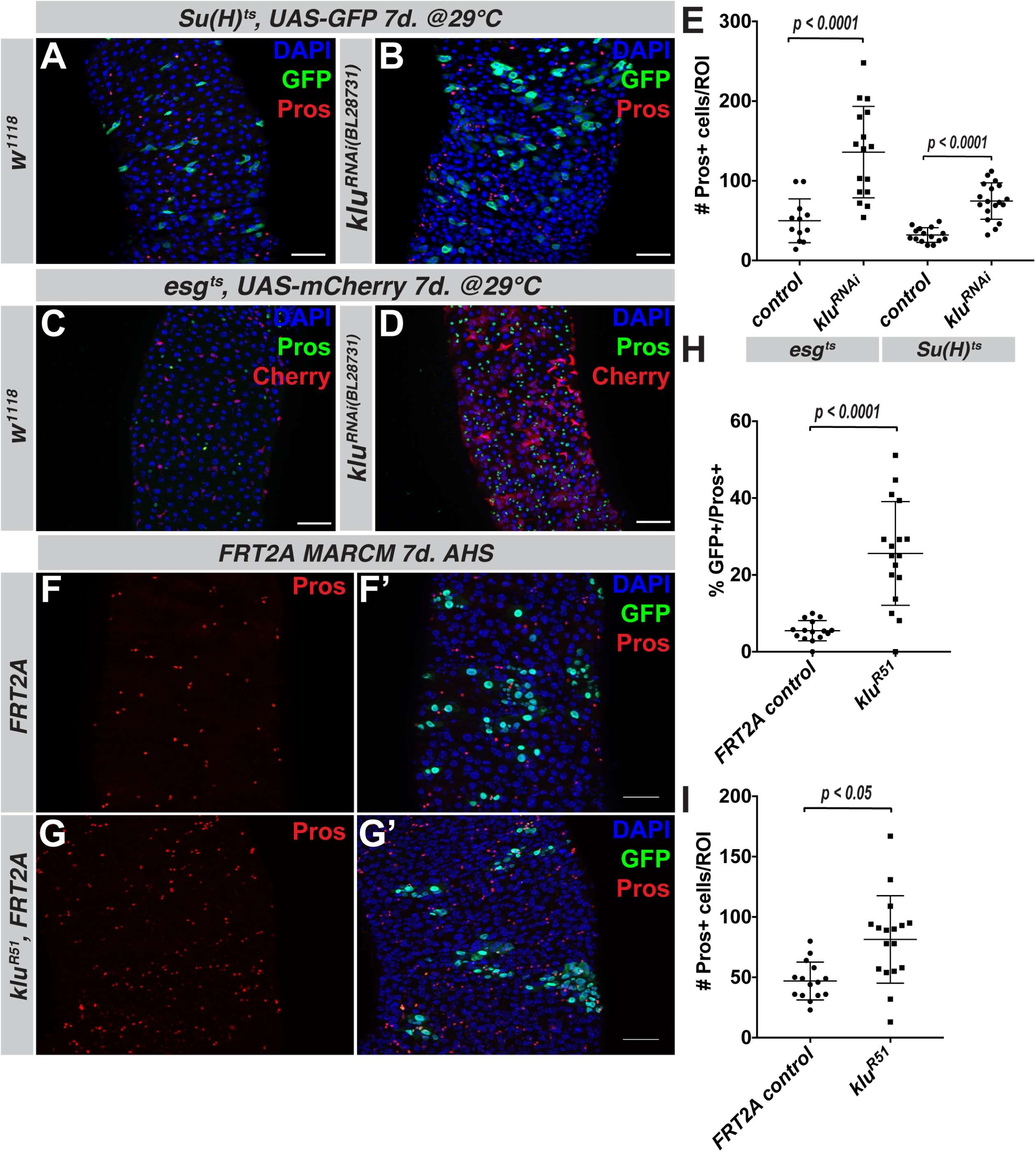
Loss of Klu leads to excess EE differentiation. **A-E**. RNAi-mediated knockdown of Klu results in an excess of Pros-positive EE-cells. Expression of *klu^RNAi^* using either the EB-specific *Su(H)GBE-Gal4^ts^* driver (Pros in red, compare A with B) or the ISC+EB driver *esg-Gal4^ts^* (Pros in green, compare C with D) results in more EE cells in the posterior midgut. **E**. EE cell quantification of the posterior midgut for the genotypes in **A-D**. Error bars represent mean +/− S.D. Significance was calculated using Student’s t-test with Welch’s correction. Number of midguts *n* = 15 (control *w^1118^*) and *n* = 18 (*klu^RNAi^*) for experiments with the *Su(H)GBE-Gal4^ts^* driver and *n* = 12 (control *w^1118^*) and *n* = 16 (*klu^RNAi^*) for *esg-Gal4^ts^* **F-I**. Clonal analysis of the *klu^R51^* null mutant allele using the MARCM technique. *klu^R51^* mutant clones have more EE cells compared to FRT2A control clones (compare F and G). **H-I**. Quantification of the number of Pros-positive EEs/clone (H) and the total number of Pros-positive EEs/ROI for the genotypes in **F-G**. *n* = 15 guts (FRT2A control) and *n* = 17 guts (*klu^R51^*). Error bars represent mean +/− S.D. Significance was calculated using Student’s t-test with Welch’s correction.

In summary, these results indicate that loss of Klu function alters the EE-to-EC ratio, but does not impair EC differentiation.

### Ectopic Klu expression inhibits stem cell differentiation and impairs Delta-Notch signaling

We hypothesized that constitutive Klu expression would impair EE differentiation. To test this, we used the *esg-F/O* system to express full-length Klu in ISC-derived clones. Wild-type *esg-F/O* clones take up most of the posterior midgut 2 weeks after induction, containing a mixture of ECs and EEs (Figure 3A). In contrast, clones expressing full-length Klu did not exhibit any hallmarks of differentiation into either EEs or ECs, but contained only a few small, round cells (Figure 3B). Klu is thought to act mainly as a repressor of transcription, based on studies in neuroblast stem cells and in sensory organ precursor (SOP) cells in the developing wing ^20,24,28^. To ask whether this repressor function of Klu would elicit the phenotypes observed, we expressed the zinc-finger DNA-binding domain of Klu fused to either a VP16 activation domain (Klu-VP16) or fused to the repressor domain from Engrailed (Klu-ERD) ^28^. Whereas differentiation still occurred in clones expressing the activating Klu-VP16, differentiation was significantly impaired in clones expressing the repressing Klu-ERD, confirming that transcriptional repression of genes regulated by Klu is sufficient to impair differentiation (Figure 3C-D, quantification in E).

**Figure 3.**
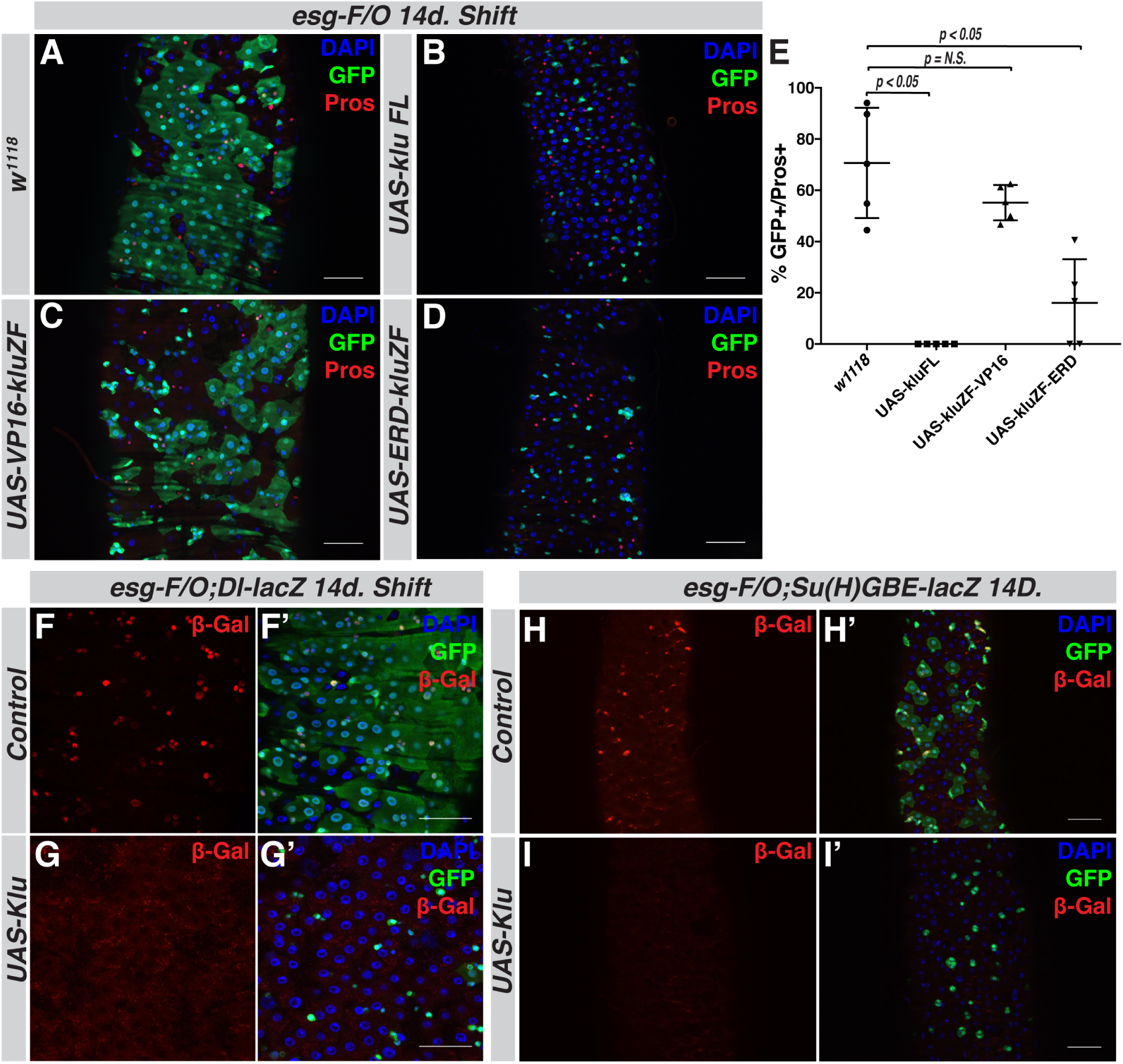
Klu overactivation results in a loss of intestinal stem cell differentiation and Delta-Notch signaling. **A-D**. Clonal expression of different Klu isoforms using the esg-FlipOut (*esg-F/O*) system to generate ISC clones. **A**. Control *esg-FO* clones grow to occupy most of the posterior midgut 2 weeks after clonal induction. **B-D**. Clones expressing either full-length Klu (*UAS-kluFL*, **B**) or the Klu zinc-finger DNA-binding domain fused to the Engrailed repressor domain (*UAS-ERD-kluZF*, **D**) resulted in a block of differentiation. This was not observed when expressing the Klu zinc-finger DNA-binding domain fused to the VP16 transcriptional activator domain (*UAS-VP16-kluZF*, **C**). **E**. Quantification of genotypes in **A-D**. Error bars represent mean +/− S.D. Significance was calculated using Student’s t-test with Welch’s correction. *n* = 5 for each genotype. **F-I**. Klu expression in ISC clones leads to a loss of Notch signaling activity in ISC-EB pairs. **F.** Control clones always contain 1 or more ISCs that are positive for the Notch ligand Delta (*Dl-lacZ*, red). **G**. *Dl-lacZ* staining is absent from clones expressing *UAS-kluFL*. **H**. Control clones have EB cells that are marked by presence of the Notch activity reporter *Su(H)-GBE-lacZ* (red). **I**. Notch transcriptional activity is absent from *UAS-kluFL*-expressing *esg-F/O* clones.

Together with the temporally restricted endogenous expression pattern of Klu, these data indicate that Klu is acting in early EBs to restrict EE differentiation, but that it has to be downregulated and that its sustained expression keeps cells in an undifferentiated state. To specify this state, we combined *UAS-Klu* with the ISC-marker *Dl-lacZ* and the EB-marker *Su(H)GBE-lacZ*. Interestingly, *esg-F/O* clones expressing *UAS-klu* did not stain positive for either *Dl-lacZ* or *Su(H)GBE-lacZ*. This suggests that Klu expression promotes an exit from the (Dl^+^) stem cell state, but also interferes with transcriptional programs induced by Delta-Notch signaling in EBs. To further investigate the interaction between Notch signaling and Klu activity in ISC differentiation, we performed epistasis experiments: the formation of large tumors consisting of proliferating ISCs and Pros-positive EEs in N loss of function conditions could be prevented by expression of Klu (*UAS-klu* expression in *N^RNAi^ esg-F/O* clones), which resulted in a complete block of both cell proliferation and EE differentiation (Figure S2). This suggests that Klu acts downstream of Notch in EC differentiation, but additionally acts as a potent inhibitor of cell proliferation. We further combined expression of Klu (*UAS-Klu*) with expression of the oncogenic *Ras^V12^* variant (*UAS-Ras^V12^*) in *esg-F/O* clones. Whereas *esg-F/O> Ras^V12^* clones occupy the entire posterior midgut 2 days after induction and contribute to a significant loss of viability of the animal, co-expression of *UAS-klu* markedly reduced clonal size at this timepoint and rescued viability (Figure S3).

Altogether, our results indicate that the N-mediated transient expression of Klu in EBs is critical to restrict lineage commitment to the EC fate and inhibit proliferation, but that normal differentiation can only proceed once Klu is downregulated. To test this hypothesis, and to better understand the mechanism by which transient Klu expression controls EB cell fate, we decided to explore the progenitor cell-specific transcriptional program downstream of Klu.

### Transcriptome profiling supports a role for Klu in regulating Notch signaling and EE fate repression

To gain a comprehensive overview of the genes controlled by Klu in the intestine, we performed RNA-Sequencing on biological triplicate populations of FACS sorted Esg-positive progenitor cells in which we expressed either a *klu^RNAi^* construct or *UAS-klu* (see Figure 4A, Supplementary Figure S5 and Methods for details). To perform these studies, we used an approach previously described in ^29^. We first confirmed that the transcriptome of sorted cells indeed reflects the excess EE differentiation phenotype seen in *klu^RNAi^* animals by performing qRT-PCR for *prospero* (*pros*) and *scute* (*sc*). The EE marker *pros* is upregulated 5-fold upon *klu^RNAi^* (Figure 4B). The proneural transcription factor Scute (*sc*) is necessary and sufficient for EE generation in the *Drosophila* midgut ^14,30,31^ and many upstream factors impinge on the expression of Sc to regulate EE differentiation ^32^. mRNA levels of *sc* increase ~2.5-fold upon *klu^RNAi^* animals and *UAS-klu* expression completely abolishes *sc* mRNA expression in stem-progenitor cells (Figure 4B).

**Figure 4.**
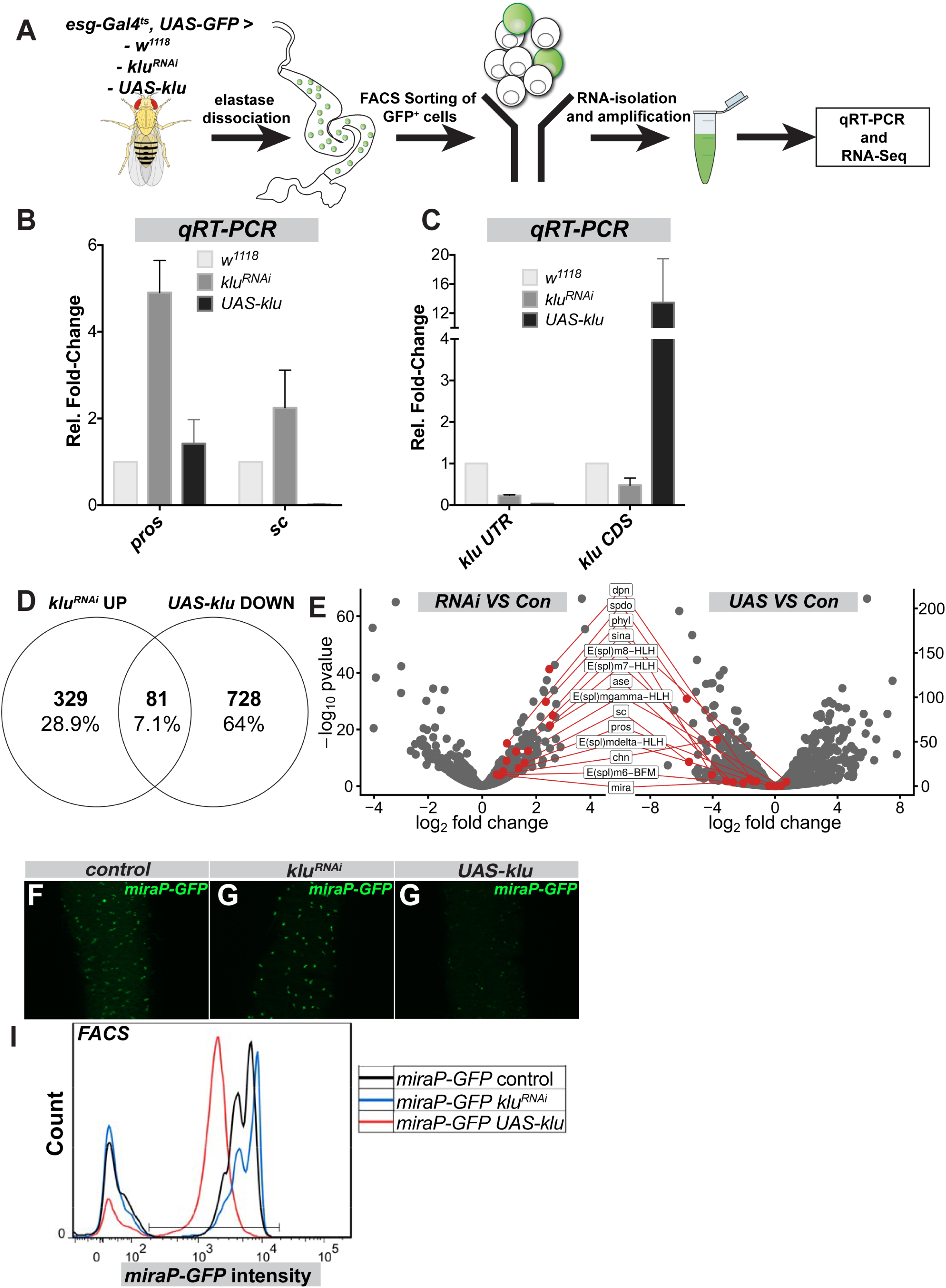
Transcriptome profiling indicates that Klu acts to repress Notch signaling output and genes critical to EE differentiation. **A**. Overview of the experiment: *esg^ts^* GFP^+^ cells either expressing *klu^RNAi^* or *UAS-kluFL* were sorted in triplicate and their transcriptome was compared to control *esg^ts^* GFP cells (see Methods for more details). **B**. qRT-PCR analysis of sorted cells for Klu and the critical EE fate regulators Scute (*sc*) and Prospero (*pros*). **C**. qRT-PCR analysis of *klu* mRNA expression with a primer pair that targets the endogenous 3’ UTR coding sequence (*klu UTR*) and a primer pair that targets the coding region only (*klu CDS*). **D**. Overlap of significantly upregulated genes in *klu^RNAi^* and significantly downregulated genes in *UAS-kluFL*. **E**. Volcano plots comparing expression of a selection of genes from the overlap of 81 genes shown in (D). Genes upregulated in the *klu^RNAi^* VS control set (left) are downregulated in the *UAS-klu* VS control set (right). **F-I**. Klu represses Da-dependent *miraP-GFP* expression in ISC and EB. Control *miraP-GFP* expression (F) is high in ISCs and EBs and slightly increased in *klu^RNAi^* midguts (G). *UAS-kluFL* expression in reduced levels of *miraP-GFP* (H). **I**. GFP-intensity of of *miraP-GFP*-positive cells for the genotypes in (F-H) by FACS. *n* = 50 midguts per genotype.

In addition, we checked the mRNA levels for Klu to verify knockdown and overexpression efficiency. As expected, we see a 70% reduction in mRNA levels upon *klu^RNAi^*. Surprisingly, we found that the expression of *klu* mRNA in *UAS-klu*-expressing progenitor cells was almost completely abolished compared to control (Figure 4C). This was contrary to our expectation of detecting increased expression upon Gal4/UAS-mediated overexpression, but was explained by the fact that the *UAS-klu* construct does not carry the endogenous *klu* 3’UTR, which our primers targeted. Accordingly, primers that target the coding region of *klu* (*klu CDS*) readily detect a ~12-fold upregulation of *klu* transcript. Hence, while transgenic Klu was induced as expected, the endogenous Klu gene was repressed, indicating that Klu may repress its own expression. The notion of a negative autoregulatory loop was confirmed in our RNA-seq data, as we saw increased reads in the CDS of the gene, and no reads in the 3’ UTR (Figure S5).

Comparing the transcriptomes of wild-type progenitors with the experimental samples, we found 410 genes significantly upregulated in *klu^RNAi^* and 809 genes significantly dowregulated in *UAS-klu*-expressing *esg^+^* cells (*Padj* < 0.05, log_2_FC > 0.5 or < −0.5). Given the role of Klu as a repressor of transcription, we focused our RNA-Seq data analysis on genes that would be upregulated in the absence of Klu, but downregulated upon *UAS-klu* expression in Esg-positive stem-progenitor cells (Figure 4D). In this category of 81 genes, many genes involved in the regulation of Notch signaling (the HES/*E(Spl)-* complex genes *m6, m7, m8*, and the HES-like transcription factor Deadpan), as well as several previously described regulators of EE differentiation (encoding the proneural proteins Asense (*ase*), Scute (*sc*), and the adaptor protein Phyllopod, *phyl*)) could be identified (Figure 4E). Additional *E(Spl*) genes (*E(Spl)-mδ* and *E(Spl)-mγ*) were significantly upregulated in *klu^RNAi^* samples, but did not significantly change in *UAS-klu* samples (Figure 4E). *E(Spl)-*genes are a group of genes activated by Notch that mediate its downstream transcriptional response ^33^. *Phyl*, in turn, acts to destabilize Tramtrack (*ttk*), a strong repressor of the *achaete-scute* complex genes *scute* and *asense*, and loss of which leads to a dramatic increase in EE numbers ^32–34^. Reciprocally, loss of *phyl* stabilizes Ttk and results in a complete loss of EE cells in the midgut ^35^. The induction of *phyl* in *klu* loss of function conditions thus explains the increase in EEs.

We also found that the expression of the transcription factor Charlatan (*chn*) is significantly downregulated by *UAS-klu* expression. Chn is a transcription factor that positively regulates the proneural genes Achaete and Scute and loss of Chn in the midgut leads to proliferation and differentiation defects in the stem-progenitor compartment ^36–38^. Hence, Klu represses the expression of several genes that have reported roles in EE differentiation.

Our transcriptome data also revealed transcriptome changes downstream of Klu that may explain the Klu-induced exit from the stem cell state: stem cell maintenance is dependent on the Class I bHLH-family member Daughterless (Da)/E47-like, since loss of *da* results in loss of ISC fate and differentiation into enterocytes ^31^. The gene *miranda (mira*) is a Da/proneural target gene that is also highly expressed in ISCs ^31,38^. Proneural factors such as Ase and Sc require Da to dimerize and regulate transcription ^39^. We found that loss of Klu results in a slight but significant upregulation of *mira* mRNA, whereas Klu overexpression results in a 2.3-fold downregulation (Figure 4E). To confirm this, we used the *mira-Promoter-GFP (mira-GFP*) line and combined this with *klu^RNAi^* and *UAS-klu*. Confocal microscopy and FACS-sorting of cells expressing either *klu^RNAi^* and *UAS-klu* confirmed that *UAS-klu* expression could reduce *mira-GFP* levels whereas a slight induction is seen in *klu^RNAi^* cells (Figure 4F-I).

Taken together, we conclude that Klu has three main functions in EBs: 1) To turn off the Notch transcriptional response by repressing E(Spl)-gene expression, 2) to promote the exit from the stem cell state by repressing genes like *mira*, and 3) To repress aberrant EE differentiation in the EB by repressing proneural gene activation. By responding to N activity and repressing its own expression, Klu further serves as a ‘timer’ for N-mediated transcriptional responses in EBs.

### Klu acts upstream of the transcription factor Scute in EE differentiation

Our genetic and transcriptional profiling experiments suggest that Klu likely acts downstream of Notch, but upstream of the proneural genes Ase and Sc in repressing EE differentiation (Figure 3, Figure 4, Figure S2). Scute plays a critical role in a transcriptional loop that regulates both ISC proliferation and the initiation of EE differentiation ^16^. Our data indicate that Klu can act to inhibit both proliferation and EE differentiation by affecting factors that genetically act upstream of Scute (Figure 3 and Figure 4). Therefore, we performed epistasis experiments with Klu and Sc to confirm this hypothesis. We generated *esg-F/O* clones that express *klu^RNAi^* in the presence or absence of *sc^RNAi^*. Clones expressing *klu^RNAi^* contained more EE cells compared to control clones (Figure 5A-B), whereas clones expressing *sc^RNAi^* are almost completely devoid of EE cells (Figure 5C). The combination of *klu^RNAi^* and *sc^RNAi^* also resulted in clones with little or no EE differentiation, as quantified by the number of Pros-positive cells (Figure 5D, quantification in 5E). This suggests that the appearance of extra EE cells in *klu^RNAi^-*expressing clones depends on Scute. To confirm that Scute would act downstream of Klu in determining EE fate, we combined overexpression of Scute and Klu. Clonal expression of Scute using the *esg-F/O* system resulted in clones consisting almost entirely of Pros-positive EE cells whereas clones expressing *UAS-klu* are completely devoid of EE cells (Figure 5F, Figure S6A-D). Co-expression of Klu and Scute leads to a marked reduction in clone size (Figure S6F) but EE differentiation was observed in a large fraction of the clones, although the percentage of differentiated cells is reduced compared to *UAS-Sc* alone (Figure 5F). We conclude that Scute can still induce EE differentiation, even in Klu gain-of-function conditions.

**Figure 5.**
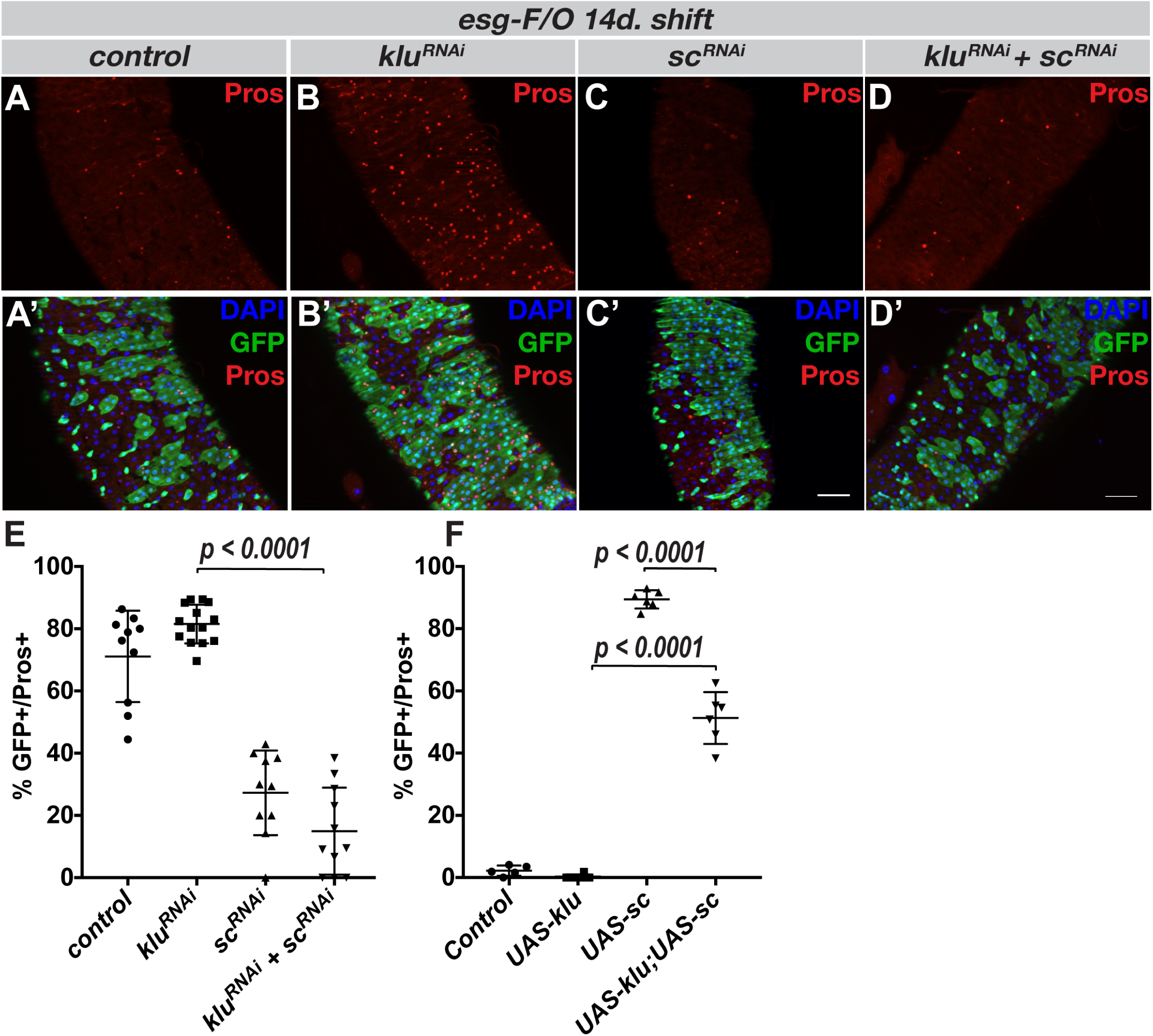
Scute acts downstream of Klu in the induction of entero-endocrine differentiation. **A-D**. Expression of *klu^RNAi^* lead to increased EE differentiation in clones 14 days after clonal induction, marked by increased numbers of Pros^+^-cells (red) (A-B, quantification in E-F). *sc^RNAi^* clones showed almost no EE differentiation (C). Similarly, the combination of *klu^RNAi^* with *sc^RNAi^* resulted in clones lacking EE differentiation (D). **E** Quantification of GFP^+^/Pros^+^ double-positive cells/clone of the genotypes in (A-D). **F**. Quantification of GFP^+^/Pros^+^ double-positive cells/clone of *esg-F/O* clones expressing either *UAS-sc*, *UAS-klu* or the combination. See Figure S6 for images. Error bars represent mean +/− S.D. Significance was calculated using Student’s t-test with Welch’s correction. For E: *n* = 10 for control and *sc^RNAi^*, *n* = 14 for *klu^RNAi^*, and *n*=12 for *sc^RNAi^;klu^RNAi^*. For F: *n* = 5 for control, *n* = 6 for *UAS-klu, UAS-sc* and *UAS-klu;UAS-sc*.

We observed an increase in the number of Pros-pH3 double-positive cells in UAS-Sc compared to control, likely representing the EE-progenitor cells (EEp) undergoing a final round of division ^16^. Strikingly, this percentage increased in *esg-F/O > UAS-klu+UAS-sc* clones (Figure S6E). Since the clonal size in this genotype is no larger than in *esg-F/O>UAS-klu* single overexpression clones (Figure S6F), indicating that these cells might be arrested in mitosis. This suggests that although Klu expression cannot completely repress *UAS-Sc*-induced EE differentiation, the effect of Klu on cell cycle progression interferes with the proliferation-inducing capacity of Scute.

### Klu directly represses several genes that regulate EE fate, Notch signaling and the cell cycle

To identify genes that are directly regulated by Klu, we performed targeted DamID of Klu in esg-positive stem-progenitor cells ^40^. We used the DamID-seq pipeline ^41^, see Methods) to identify 1667 genes that had 1 or more Klu binding peak(s) within 2 KB of their gene body in all 3 replicates. Using 2 published position weight matrices for Klu-binding ^42^, we could establish that 692 of the 1667 genes (41.5%) had 1 or more Klu-binding motif(s) present in their binding peaks. We considered these peaks as high-confidence Klu-bound sites. Our transcriptomics data on Klu indicated that Klu controls many genes involved in Notch signaling, EE differentiation and cell cycle regulation. We identified a cluster of binding sites at the centrosomal end of the *E(Spl)-*locus around the *E(Spl)-mδ* and *E(Spl)-mγ* genes (Figure 6E). Since our RNA-Seq data showed that many of the *E(Spl)-*genes change expression in both *klu^RNAi^* and *UAS-klu* conditions (Figure 4E), this suggests that Klu could possibly regulate the expression of multiple members of the *E(Spl*)-complex through this binding peak at the centrosomal end of the *E(Spl*)-locus. Furthermore, we identified a Klu-Dam binding peak at the *klu* locus, supporting our hypothesis that Klu acts in an autoregulatory loop by negatively regulating its own expression (Figure 6A). Previous work has shown that Scute and the *E(Spl*)-complex member *E(Spl)m8-HLH* act in a regulatory loop to generate an EE precursor directly from the ISC ^16^. Since our results indicate that Scute is upregulated upon loss of Klu and acts epistatically to Klu in EE formation, we first looked for Klu binding in and around the *scute* locus. We did not find significant binding of Klu-Dam around any of the genes in the Achaete/Scute complex. However, we did identify a DamID peak around the *sina* and *sinah* loci (Figure 6B). Together with the adaptor protein Phyllopod, the Sina and Sinah E3-ubiquitin ligases are able to degrade the transcriptional repressor Tramtrack (*ttk*), which represses EE fate ^32,35^. *sina* transcript levels are upregulated 2.2-fold upon *klu* RNAi and *phyl* levels are upregulated 8-fold as well as downregulated 15-fold upon *UAS-klu* expression (Figure 4E). Hence, we propose that Klu represses EE-fate determination in EBs upstream of Scute by stabilizing Tramtrack, since Klu directly represses the members of the E3-ligase complex Sina, Sinah and (indirectly) Phyl that can normally target Ttk for destruction.

**Figure 6.**
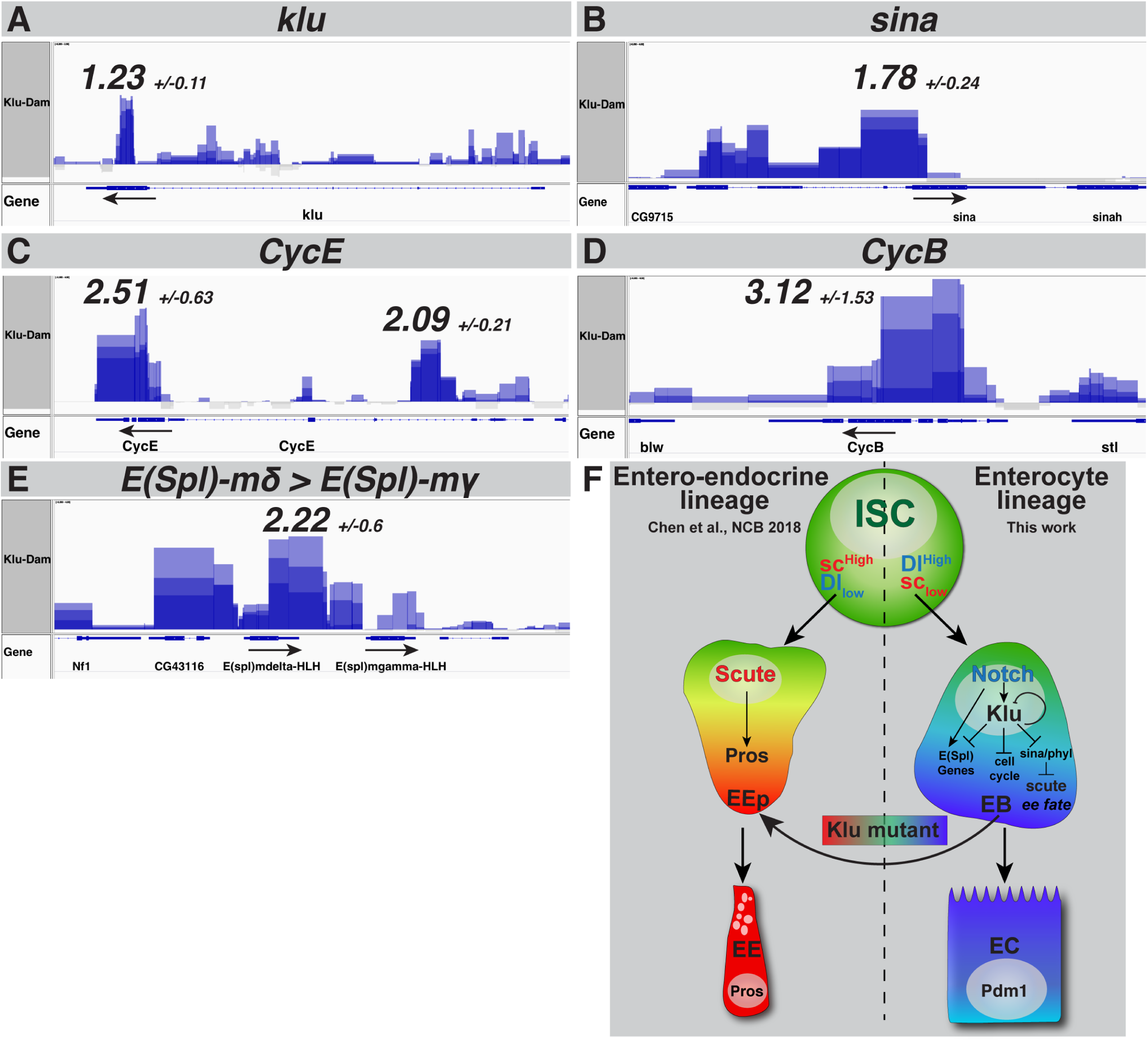
Klu directly represses genes involved in Notch signaling, cell cycle and EE differentiation. **A-E**. Klu-Dam binding tracks (in triplicate) for the *klu* locus (A), *sina* locus (B), *Cyclin E* (C) and *Cyclin B* (D) as well as the E(Spl)-complex genes *E(Spl)-mδ* and *E(Spl)-mγ* (E). Tracks are displayed using Integrated Genome Viewer (IGV) as overlayed tracks of the triplicate Klu-Dam VS Dam-only control comparisons. Arrows indicate direction of transcription. Numbers indicate the average maximum height of the peak (log_2_FC of Klu-Dam over Dam-only control) for each of the 3 replicates. **F**. Model of Klu function in lineage differentiation in the intestine. Klu expression is activated by Delta-Notch signaling in the EB cell, together with other members of the Hairy/Enhancer of Split family of Notch target genes. Klu accumulation in EBs results in a subsequent repression of these target genes, including the repression of its own expression. Additionally, Klu acts as a safeguard to repress erroneous EE differentiation in the enteroblast-enterocyte lineage by indirectly repressing the accumulation of proneural genes such as Asense and Scute through inhibition of the E3-complex members Phyllopod (*phyl*) and Seven-in Absentia (*sina*) that repress the accumulation of Scute and thereby inhibit EE fate. Finally, Klu acts in the regulation of the cell cycle in EB cells, as the cell remodels its cell cycle from a mitotic to an endocycle.

In addition to genes involved in Notch signaling and EE-specification, we find evidence for direct repression of critical cell cycle regulators by Klu. We find Klu binding peaks at both the Cyclin B (*CycB*) and Cyclin E (*CycE*) loci (Figure 6C and D), two Cyclins that are essential for G1-S and G2-M progression respectively. CycE is also upregulated upon *klu* RNAi expression. Notch activation is essential for the mitotic-to-endocycle switch in follicle cells of the Drosophila ovary, and polyploidization is a critical step in the normal process of EB-to-EC differentiation ^43,44^. We propose that Klu plays a role in remodeling the cell cycle in response to Notch activation by directly repressing 2 critical cell cycle regulators. Furthermore, this explains how Klu acts as a potent suppressor of cell proliferation, even in combination of the strongly oncogenic RasV12 overexpression (Figure 3 and Figure S3).

Altogether, our data suggest a model (Figure 6G) in which Klu acts as a Notch effector in the EB that acts to restrict the duration of the Notch transcriptional response (through negative regulation of the E(Spl)-complex members and Klu itself). Second, Klu prevents activation of the Scute-E(Spl)m8 transcriptional circuit that triggers EE differentiation. Finally, we find evidence that Klu can bind and repress critical cell cycle regulators such as Cyclin B and Cyclin E, likely promoting the switch from a mitotic to an endoreplicating cell cycle in differentiating ECs.

## Discussion

Our work identified a mechanism by which lineage decisions are cemented through the coordinated repression of alternative fates and of cell proliferation in somatic stem cell daughter cells. N-induced expression of Klu in EBs is necessary to repress EE fates in EBs, but also to temporally restrict N target gene expression. Hence, its own expression has to be self-regulated to allow differentiation to ECs to proceed. We find that Klu represses several genes that are critical for EE differentiation; most notably genes that influence the accumulation of the transcription factor Scute. Transient expression of Scute is necessary and sufficient for EE differentiation and the transient nature of Scute activation is accomplished by a double-negative feedback loop between Scute and E(Spl)m8 ^16^. The expression of Klu results in the repression of both transcription factors in EBs, inactivating the transcriptional circuit that governs EE differentiation (Figure 6F). Previous work on Klu has shown that Klu is directly regulated by Su(H) and acts as a Notch effector in hemocyte differentiation ^45^. In the ISC lineage, we find that overexpression of Klu results in the loss of Notch signaling activity in progenitor cell clones, and that Klu is able to repress several Notch effector genes (such as the HES/E(Spl) family and HES/E(Spl)-like genes such as Deadpan). We thus propose that Klu acts in a negative feedback loop downstream of Notch signaling to ensure that Notch effector gene activity is transient in EBs, mirroring the transient nature of EE specification by Scute and E(Spl)m8.

Loss of WT1 in the mouse kidney results in glomerulosclerosis and is accompanied by ectopic expression of HES/E(Spl) family genes ^46^ and in zebrafish kidney podocytes Notch expression induces *Wt1* transcription, while the Notch intracellular domain (NICD) and WT1 synergistically promote transcription at the promoter of the HES/E(Spl) family gene Hey1 ^47^. This suggests that the negative feedback between Notch and its effector Klu/WT1 might be conserved between species, even though conservation at the sequence level between these transcription factors is low.

Our data also support a role for Klu for regulating cell cycle progression. Overexpression of Klu results in a strong block in cell proliferation in *N^RNAi^* or oncogenic *Ras^V12^*-induced tumors and our DamID binding studies suggest that Klu can directly regulate Cyclins B and E. This is in stark contrast to its role in the neuroblast stem cell lineage, where overexpression of Klu leads to a strong overproliferation of immature neural progenitor cells and the resulting formation of brain tumors that can be transplanted to distant regions of the body and continue to grow ^20,21^. However, this likely reflects the different role for Notch in the NB lineage, where continuous activation of Notch also leads to INP overproliferation and tumor formation. Thus, the role of Klu in promoting either lineage differentiation or stem-progenitor cell proliferation seems to be context-dependent. This is reminiscent of the context-specific rules for Notch as either a tumor suppressor or an oncogene in mammalian adult stem cell lineages. For instance, activating Notch1 mutations are the main driver of T-cell acute lymphoblastic leukemia ^48^, whereas Notch-inactivating mutations are found in many epithelial-derived solid tumor types such as head-and-neck or skin carcinomas (see ^49^ and references therein). Similarly, *Wt1* was initially identified as a tumor-suppressor gene mutated in the rare pediatric kidney cancer Wilms’ Tumor ^50^. Recently, WT1 was also identified as a strong inhibitor of proliferation in a genome-wide gain-of-function screen for tumor suppressors and oncogenes ^51^. However, expression of WT1 was found to be elevated in many solid tumors and in acute myeloid leukemia ^52,53^. Thus, similar to Notch, the role of WT1 in tumorigenesis seems to be highly context-dependent. Intriguingly, in both nephric and hematopoietic lineages WT1 is often transiently expressed in committed progenitor cells, similar to the expression of Klu in the EB progenitors, raising the possibility that to fully understand the role of WT1-like proteins in tumorigenesis, cell lineage relationships, as well as cell proliferation and differentiation events in tumors need to be taken into account.

Critically, our work highlights the role for transient transcriptional ‘rewiring’ events during cell specification in somatic stem cell lineages. Such events seem to be required to ensure lineage commitment downstream of initial symmetry breaking signals like Notch, and ensure progression of cell differentiation into a defined lineage. As such, it can be expected that similar transcriptional feedback loops need to be reversed for cells to undergo de-differentiation into stem cells in mammalian regenerating tissues. We propose that understanding their makeup and regulation will significantly advance efforts to control tissue repair and regeneration in mammals, including humans.

## Methods

### Fly strains and husbandry

The following strains were obtained from the Bloomington Stock Center: BL28731 (klu RNAi on 3rd) BL60469 (klu RNAi on 2nd), BL56535 (*UAS-klu*[*^Hto^*]), BL11651 (*Dl^05151^-lacZ*) BL26206 (*sc RNAi*), BL51672 (*UAS-sc*), BL1997 (*w*[*\**]*; P{w*[*+mW.hs*]=*FRT(w*[*hs*]*)}2A*), BL4540 (*w*[*\**]*; P{w*[*+mC*]=*UAS-FLP.D}JD2*). BL65433 (*y*[*1*] *w*[*\**]*;M{w*[*+mC*]=*hs.min(FRT.STOP1)dam}ZH-51C*) BL1672 (*w*[*1118*]*; sna*[*Sco*]*/CyO, P{ry*[*+t7.2*]=*en1}wg*[*en11*]). VDRC: v27228 (*N RNAi*). Other stocks: *klu-Gal4 UAS-GFP*, *FRT2A kluR51/Tm6B*, *hs-Flp, Tub-Gal4, UAS-GFP/Fm7;FRT2A, TubGal80ts/Tm2,Ubx* (T. Klein, Düsseldorf) *UAS-kluFL, UAS-ERD-kluZF, UAS-VP16-kluZF* (C.Y. Lee, U. Michigan) *esg-F/O* (*w; esg-Gal4, tub-Gal80ts, UAS-GFP; UAS-flp, Act>CD2>Gal4(UAS-GFP)/TM6B*), *esg^ts^* (*y,w;esg-Gal4, UAS-GFP/CyO;tub-Gal80ts/Tm3*), *Su(H)^ts^* (*w;Su(H)GBE-Gal4,UAS-CD8-GFP/CyO;tub-Gal80^ts^/TM3*), ISC-specific *esg^ts^* (Wang et al., 2014) *w;esg-GAL4,UAS-2XEYFP/CyO;Su(H)GBE-GAL80,tub-Gal80ts/TM3,Sb*, *w;esg-gal4, tub-Gal80ts, UAS-GFP/CyO,wg-lacZ;P{w*[*+mC*]=*UAS-FLP.D}JD2/Tm6B*. Stocks generated in this study: *w;Klu-Dam(ZH-51C) M4M1/CyO, P{ry*[*+t7.2*]=*en1}wg*[*en11*]

### Immunostaining and microscopy

Midgut immunostaining was performed as described in ^54^. Antibodies used include: Chicken anti-GFP (1:1000, ThermoFisher A10262), mouse anti-Prospero (MR1A, 1:50, DSHB), mouse anti-β-galactosidase (40-1a, 1:200, DSHB), rabbit anti-β-galactosidase (1:200, ThermoFisher A11132) mouse anti-Armadillo (N2 7A1, 1:20, DSHB), rabbit anti-phosphorylated Histone H3-Ser10 (pH3S10, 1:500, sc8656-R, Santa Cruz Biotechnology). Images were taken from the R5 and R4 regions of the posterior midgut on a Zeiss Apotome microscope or Zeiss LSM710 confocal at either 20X or 40X magnification. Images were captured as Z-stacks with 8-10 slices of 0.22-1.0 μm thickness. Images were converted to maximum-intensity projections in Fiji (https://fiji.sc) and quantifications were performed using the CellCounter FiJi plugin. Scale bar = 50 μm in all images, except in Figure 1A: scale bar = 25 μm. Graphing, statistical analysis and survival curves were produced in GraphPad Prism.

### Cloning and transgene generation

We used the Inducible DamID system from the Van Steensel lab to generate klu-Dam ^40^. To this end, we amplified the Klu Full-length cDNA (derived from BDGP Gold clone FI01015) using AscI and NotI-containing primers and cloned the fragment into the vector *p-attB-min.hsp70P-FRT-STOP#1-FRT-DamMyc*[*open*] (Addgene plasmid #71809). Transgenic lines were generated by Genetivision Inc. using the phiC31 integrase-mediated site-specific transgenesis system ^55^. The finished construct was injected into Bloomington stock BL24482 (*ZH-51C* attP-site on 2nd) and the resulting transgenic lines were tested by genotyping PCR. Both control (Dam-only, BL65433) and klu-Dam transgenic lines were crossed to BL1672 (*w*[*1118*]*; sna*[*Sco*]*/CyO, P{ry*[*+t7.2*]=*en1}wg*[*en11*]) before use.

### DamID

Control Dam-only (BL65433) and klu-Dam male flies were crossed to *w;esg-gal4, tub-Gal80ts, UAS-GFP/CyO,wg-lacZ;P{w*[*+mC*]=*UAS-FLP.D}JD2/Tm6B* virgins. Crosses were maintained at 18°C and progeny was shifted to 29°C for 24 hours to induce the Flp-mediated recombination of the STOP-Cassette. 30-50 midguts of Dam-only and klu-Dam were dissected in 1X PBS in 3 different batches and used for isolation of total genomic DNA. Isolation of methylated GATC-sequences and subsequent amplification was done according to the protocol published by Marshall et al. ^56^ until Step 34, from which we continued NGS library preparation using the Illumina TruSeq nano DNA kit LT. After library quality control, samples were sequenced as 50 bp single-end on an Illumina HiSeq2500.

### Midgut FACS, RNA-isolation and Sequencing

Midgut dissociation and FACS was performed as described in ^29^. UAS-expression of *UAS-klu* or *klu^RNAi^* was induced for 2 days, followed by 16 hours of *Ecc15* infection to stimulate midgut turnover. We dissected 100 midguts/genotype in triplicate and for each sample 20,000-40,000 cells were sorted into RNAse-free 1X PBS with 5 mM EDTA. RNA was isolated using the Arcturus PicoPure™ RNA Isolation Kit. Subsequently, the entire amount of isolated RNA was used as input for RNA-amplification using the Arcturus™ RiboAmp™ HS PLUS Kit. 200 ng of amplified aRNA was used as input for RNA-Seq library preparation using the TruSeq Stranded mRNA Library Prep Kit (Illumina) and samples were subsequently sequenced as 50 bp single-end on an Illumina HiSeq2500.

### Quantitative real-time PCR

Quantitative real-time PCR (qRT-PCR) was performed using aRNA from FACS-sorted esg^+^ cell populations (see above) as template. qRT-PCR was performed using the TaqMan FAM-MGB system in a 10 μl reaction on a BioRad CFX384 C1000 Touch Cycler using the following probes: klu (dm02361358 s1), pros (dm02135674 g1), sc (dm01841751 s1) and Act5C (dm02361909 s1) was used for normalization. The klu CDS primer assay was ordered as a Custom TaqMan Assay. Reactions were performed in triplicate on 3 independent biological replicates. Relative expression was quantified using the ^ΔΔ^Ct method. Data were calculated using Microsoft Excel and plotted as relative fold-changes +/− SEM in Graphpad Prism.

### RNA-Seq and DamID data analysis

The 15-21 million quality-passed reads per sample were mapped to the *D. melanogaster* reference genome (BDGP6) with TopHat2 (version 2.1.0) ^57^. Of each sample, approximately 80% of the reads was mapped to the genome. From this, 90% could be assigned to genes using FeatureCounts resulting in 11-15 million analysis-ready reads per sample ^58^.

The table of raw counts per gene/sample was analyzed with the R package DESeq2 (version 1.16.1) for differential expression ^59^. Both sample groups of interest (UAS & RNAi) were pair-wise contrasted with the control sample group (control). For each gene of each comparison, the p-value was calculated using the Wald significance test. Resulting p-values were adjusted for multiple testing with Benjamini & Hochberg correction. Genes with an adjusted p-value <0.05 are considered differentially expressed (DEGs). For DamID, we used the damid_seq pipeline ^41^to genrate binding profiles for Klu-Dam. Triplicate samples for Klu-Dam (34.9, 33.5 and 34.1 millions reads) and Dam-only control (34.7, 34.5, and 35.6 million reads) were aligned to the *Drosophila* genome (UCSC dm6). Overall aligning rate was between 86% and 91% across all samples. First, ‘gat.track.maker.pl’ script was used to build a GATC fragment file. Then the main utility ‘damidseq_pipeline’ was used to align the reads to the genome using bowtie2, bin and count reads, normalize counts, and compute log2 ratio between corresponding DamID and control Dam-only samples. The pipeline identified 1,707, 1,663, 1,681 peaks with FDR<0.01 per each replicate. To test for reproducibly we first used Marshall OJ’s damid_pipeline to identify peaks with weaker confidence (FDR< 0.1) and the ‘idr’ python package (https://github.com/nboley/idr) to identified 1,169 peaks with IDR<0.05 between replicate1 and replicate2. We used an in-house developed script to annotate peaks in proximity to genes. 1,667 genes found to be in proximity to at least one reproducible peak. To find Klu binding motifs in our reproducible peak set, we scanned for 2 different Klu PWM (described in ^42^) around reproducible peaks using the FIMO tool ^60^. Reads were visualized using IGV as overlayed triplicate Klu-Dam (log_2_FC over Dam-only) tracks.

## Acknowledgements

The authors would like to thank Thomas Klein, C.Y. Lee, Sarah Bray, Claude Desplan, Benoit Biteau, Cai Yu and Bruce Edgar for providing fly stocks and antibodies, the Bloomington *Drosophila* Stock Center, The VDRC (Vienna) and the Developmental Studies Hybridoma Bank (DSHB) for fly stocks and antibodies, and Maria Locke, Marco Groth, Philipp Koch and Karol Szafranski from the Flow Cytometry, Sequencing and Bioinformatics Core Facilities at the FLI-Leibniz Institute on Aging for expert technical assistance. This work was supported by DFG research grant number KO5594/1-1 to J.K.

## Author contributions

J.K and H.J conceived the project and designed experiments. J.K., M.B. and E.M. performed experiments and collected data. T.R-O. performed data analysis on the RNA-Seq and DamID samples. P.S-V. provided preliminary data for the study. J.K. and H.J. wrote the manuscript.

