## Supplementary Figures S1-S6 for "The WT1-like transcription factor Klumpfuss maintains lineage commitment in the intestine"

**Korzelius et al. 2018 Supplementary Figures**

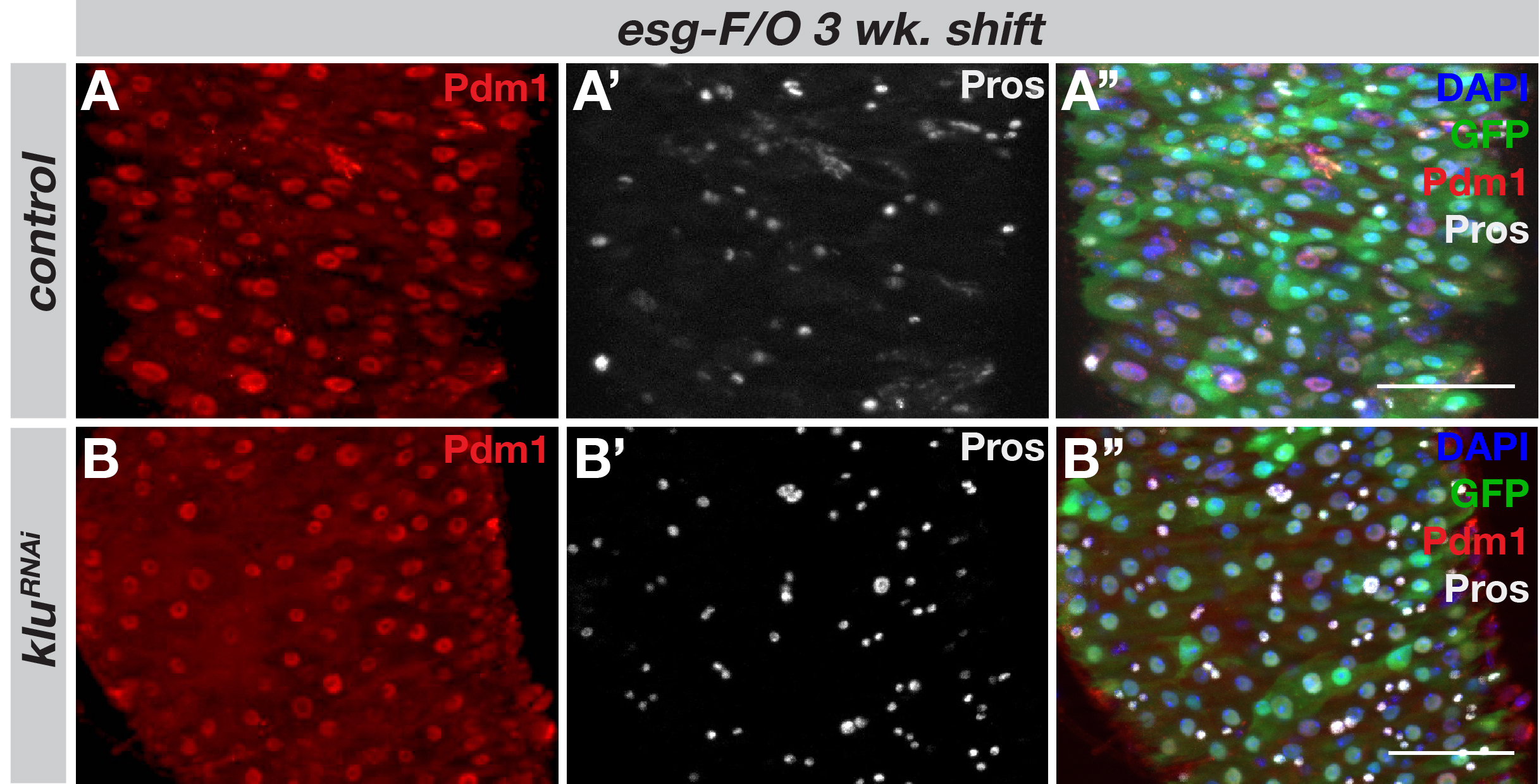

**Figure S1. Loss of Klu results in a shift in the EC-to EE ratio, but not to misdifferentiation of ECs. A-A''.** Control *esg-F/O* clones consisted of large, Pdm1-positive enterocytes (A) and Pros-positive EE cells (A') and took up most of the posterior intestine 3 weeks after clonal induction. **B-B''**. Expression of *klu^RNAi^* increased the ratio of Pros-positive cells (B'), but Pdm1-positive ECs can still form in these clones and appear similar in ploidy and Pdm1-content (B).

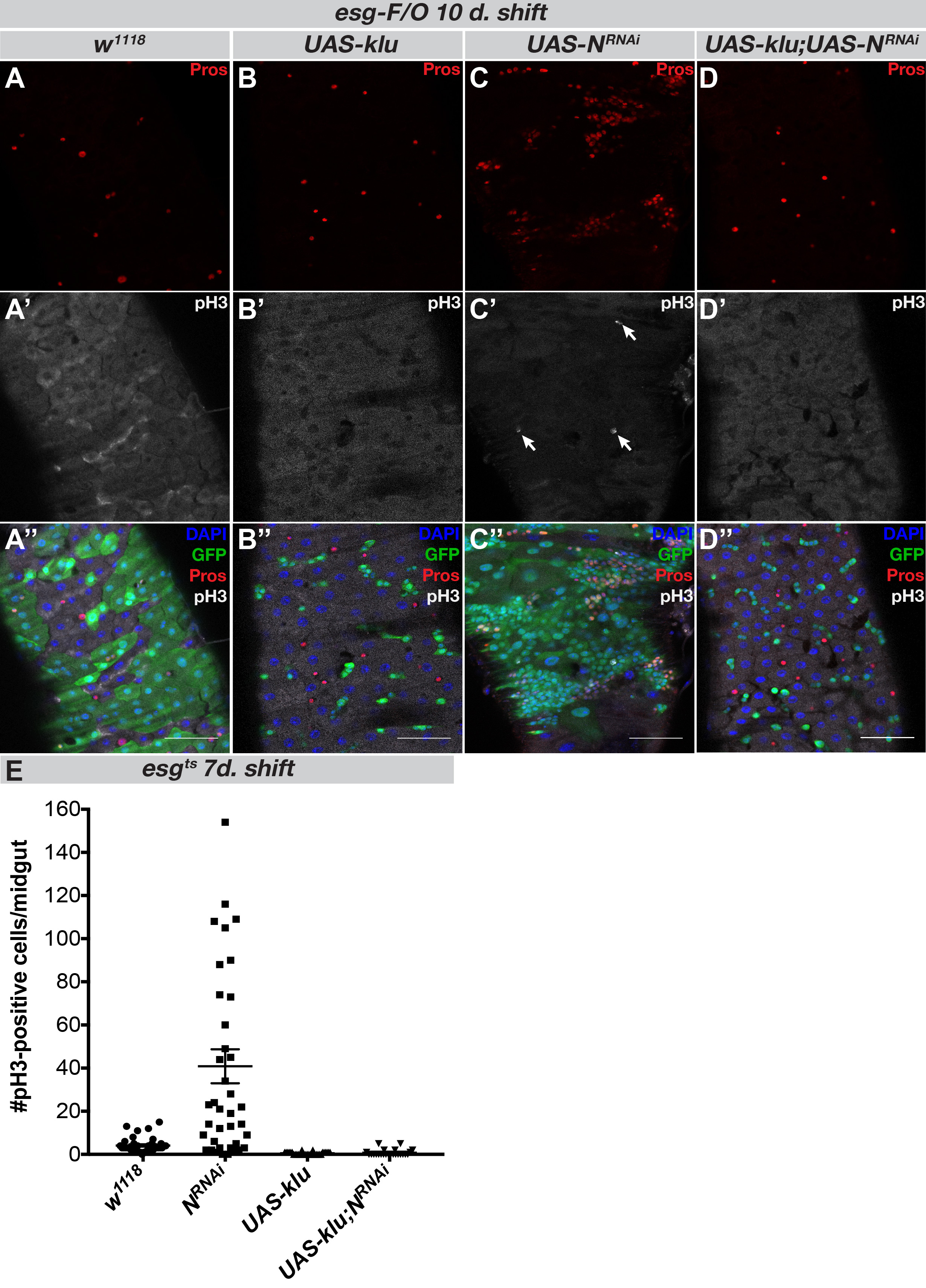

**Figure S2. Klu acts downstream of Notch in *N^RNAi^*-induced tumor formation in the intestine. A-D.** Control *esg-F/O* clones differentiated into ECs and EEs and occupied most of the posterior midgut 10 days after clonal induction (A), whereas *UAS-klu* clones fail to proliferate or to differentiate (B). Loss of Notch from *esg-F/O* clones lead to the formation of tumors consisting of mitotic ISC-like cells (arrows indicate mitoses) and Pros-positive EE cells (C). This phenotype is reversed when combining *N^RNAi^* with *UAS-klu* expression (D). **E**. The *esg^ts^* system was used to express the abovementioned constructs for 7 days and mitotic figures/midgut were counted for each genotype. The *UAS-klu;N^RNAi^* midguts had significantly less mitoses than *N^RNAi^* animals (*p<0,0001*, Mann-Whitney U-test). *n*=42 for control (crossed with *w^1118^*) *esg^ts^* animals and *N^RNAi^* animals, *n*=33 for *UAS-klu* and *n*=24 for *UAS-klu;N^RNAi^*.

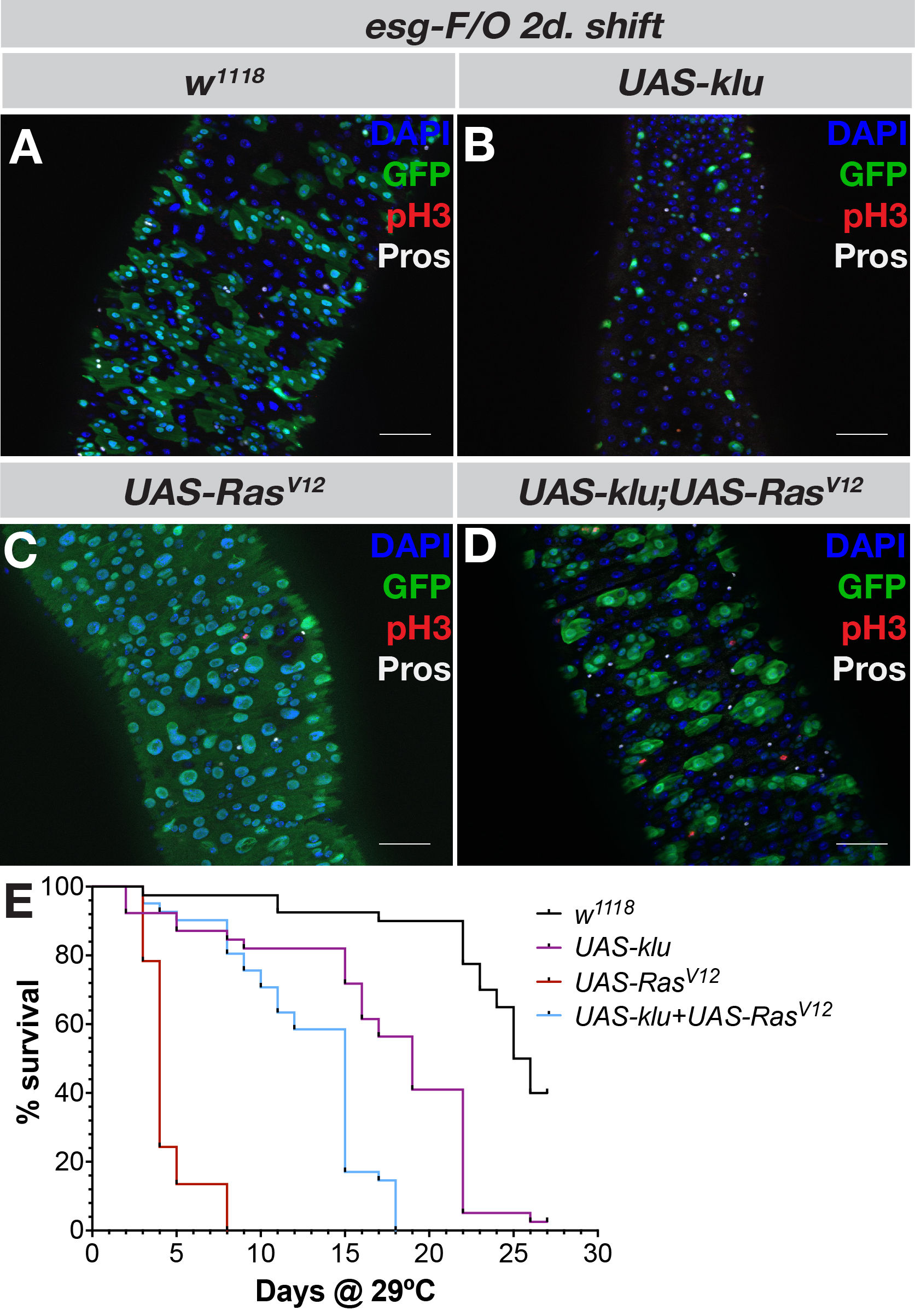

**Figure S3. Klu overactivation represses Ras^V12^-induced overgrowth.** **A-D**: After 2 days of induction, control *esg-F/O* clones (A) were mostly 1-2 cell clones, similar to *UAS-klu*-expressing clones (B). In contrast, *esg-F/O* clones expressing the oncogenic form of Ras, *UAS-Ras^V12^*, occupied the complete intestine 2 days after induction. Co-expression of *UAS-klu* with *UAS-Ras^V12^* (D) markedly reduced the Ras^V12^-induced overgrowth. **E**. Survival assay of *esg-F/O* flies at 29ºC expressing the abovementioned constructs. Co-expression of *UAS-klu* significantly extends the lifespan of *UAS-Ras^V12^* expressing flies. *n*= 40 control, *n*= 39 *UAS-klu*, *n*= 37 *UAS-Ras^V12^* and *n*=41 *UAS-Ras^V12^;UAS-klu*.

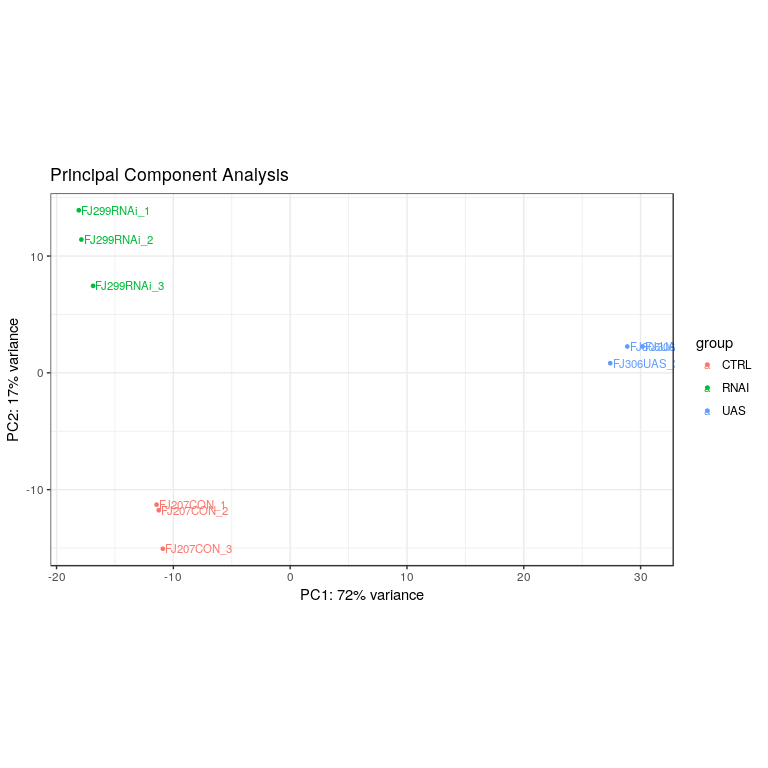

**Figure S4. PCA-analysis plot of RNA-Seq samples of control *esg^ts^* animals or animals expressing either *UAS-klu* or *klu^RNAi^* as described in Figure 4**. PCA analysis revealed a close correlation between all biological replicates in PC1 and PC2.

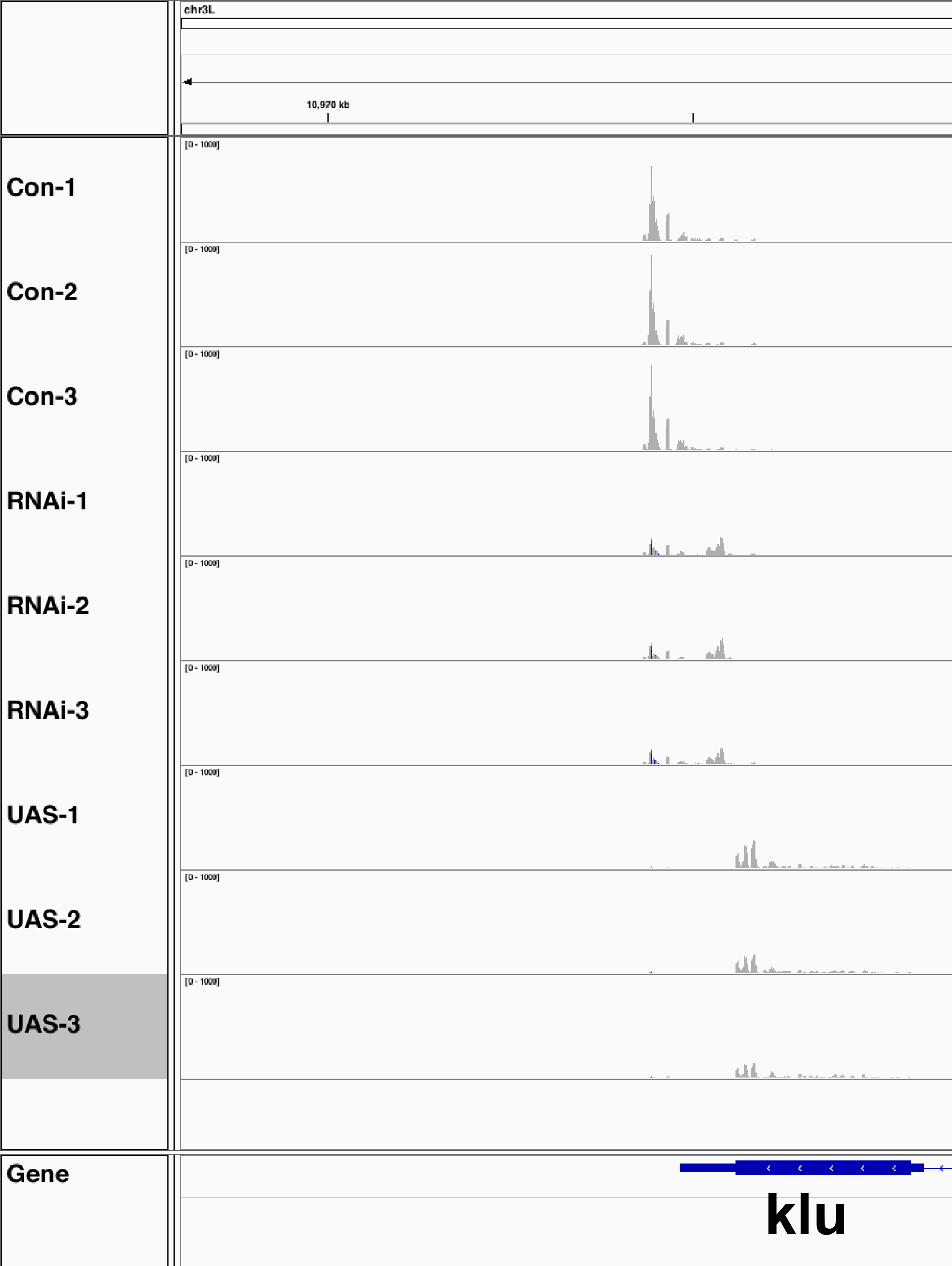

**Figure S5. IGV view of reads around the *klu* locus in all RNA-Seq samples**. The number of reads near the 3' end of the *klu* gene is high in control Esg-positive sorted cells, but reduced in samples expressing *klu^RNAi^.* *UAS-klu* samples show an absence of reads from the 3' end, but more reads along the coding region of the gene.

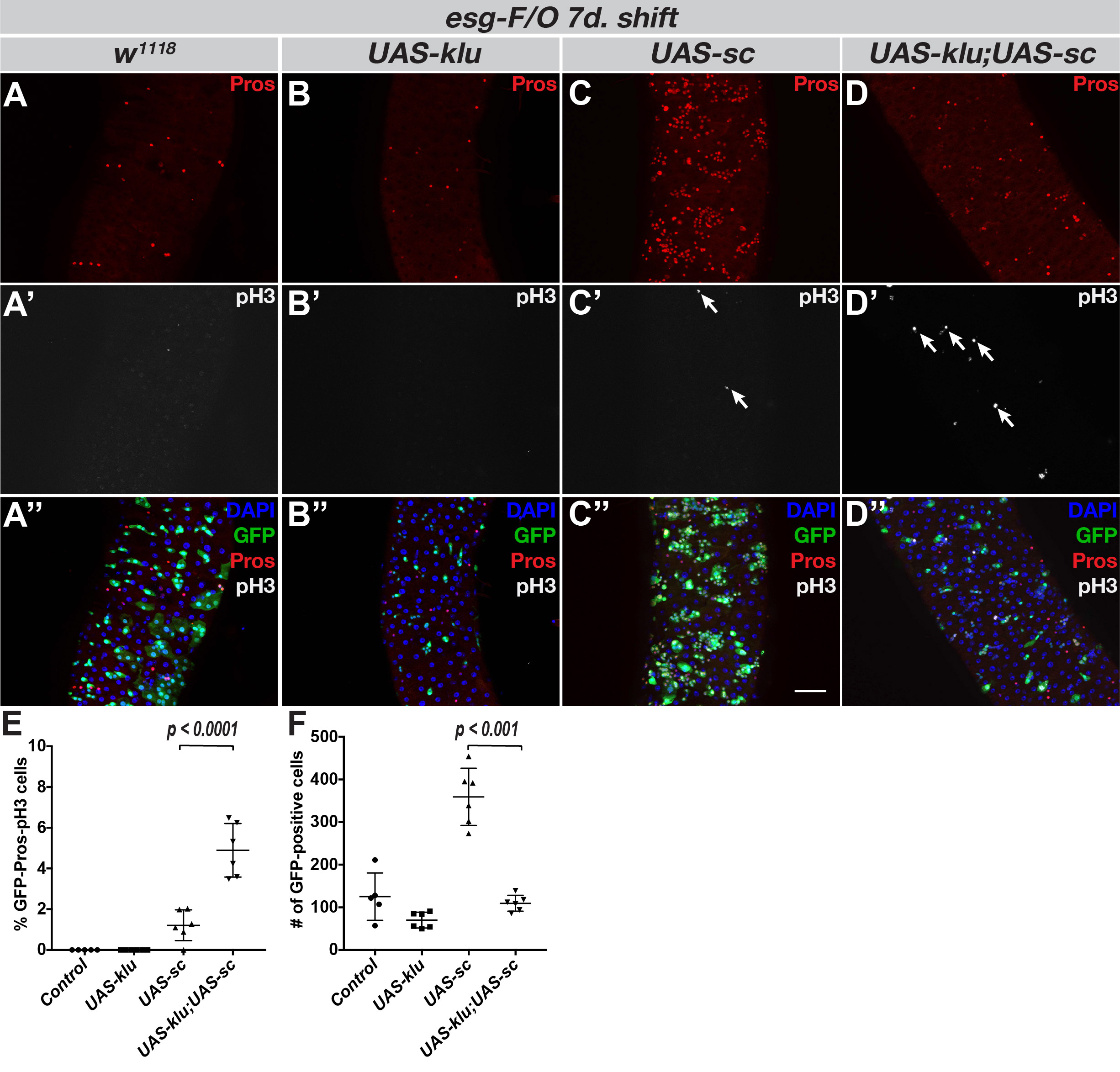

**Figure S6. Klu inhibits clonal proliferation, but not EE differentiation in *esg-F/O* clones**. **A-D**. Control *esg-F/O* clones (A) have approximately 5% EE cells, marked by Pros (red) 7 days after clonal induction. Ectopic Klu expression completely blocks differentiation (B), whereas Scute overexpression results in clones consisting almost entirely of EE cells (C). *UAS-sc* clones also show increased numbers of mitotic cells (indicated by staining for pH3S10 in white, arrows). Co-expression of Klu and Scute results in clones that still have EE differentiation, albeit at a lesser rate as clones expressing only Scute (D). These clones contain fewer cells that *UAS-sc* clones, but more pH3S10-positive cells/clone (arrows). **E**. Quantification of GFP^+^/Pros^+^/pH3^+^ triple-positive cells/clone of the genotypes in (A-D). **F**. Quantification of the total number of GFP^+^ cells/ROI of the genotypes in (A-D). Error bars represent mean +/- S.D. Significance was calculated using Student's t-test with Welch's correction. *n* = 5 for control, *n* = 6 for *UAS-klu, UAS-sc* and *UAS-klu;UAS-sc*.
